# Midzone bundles of the mammalian anaphase spindle are mechanically coupled both locally and globally

**DOI:** 10.64898/2026.04.08.716168

**Authors:** Zachary Mullin-Bernstein, Sterre van Wierst, Carly Garrison, Nathan Cho, Sophie Dumont

**Author notes:** Correspondence to Zachary Mullin-Bernstein; Sophie Dumont. Sterre van Wierst’s current affiliation: Cell Biology and Immunology, Wageningen University & Research, Wageningen, the Netherlands, Wageningen, Netherlands. Carly Garrison’s current affiliation: Vir Bio, San Francisco, CA, USA.

## Abstract

Robust chromosome segregation requires the anaphase spindle to both preserve and remodel its structure under force. How it does so remains unclear as probing mechanics during anaphase’s short lifetime is challenging. Here, we use microneedles to pull on mammalian anaphase midzone bundles and ask how they respond to and transmit force across space and time. We find that midzone bundles locally transmit force to each other in the spindle’s short axis over multiple timescales. Along the spindle’s long axis, midzone bundles globally transmit force: rather than bundles sliding apart or detaching under force, the spindle shortens. This reveals strong anchorage and resistance to outward sliding, and that spindle elongation requires all midzone bundles to elongate. This global force transmission is stronger for short-lived forces and is strictly PRC1-dependent, indicating limited mechanistic redundancy. In sum, the anaphase spindle acts as a single mechanical unit over short timescales to resist and transmit force, while remodeling over long timescales to segregate chromosomes.

**Summary:** Using microneedle manipulation, Mullin-Bernstein et al. show that microtubule bundles of the mammalian anaphase spindle are mechanically coupled, transmitting force both laterally and between spindle poles. These strong connections may help ensure coordinated and error-free chromosome segregation.

## Introduction

The spindle is the macromolecular structure responsible for orchestrating the accurate segregation of genetic material during cell division. Once the spindle builds itself and properly attaches to chromosomes, it actively separates sister chromatids in a process known as anaphase. During anaphase, the spindle must dramatically elongate and deform to move chromosomes. Yet, it must also maintain defined and robust structures, including the central spindle which is necessary for cytokinetic signaling (Jiang *et al*., 1998). Anaphase spindle elongation can help resolve erroneously attached chromosomes (Cimini *et al*., 2004; Scholey *et al*., 2016), while failures of anaphase functioning can cause aneuploidy (Gordon *et al*., 2012; Lens and Medema, 2019). How the anaphase spindle can be so structurally dynamic while remaining functionally robust remains an open question. Specifically, our understanding of the mammalian anaphase spindle’s emergent mechanics remains understudied due to the challenges of probing mechanics in live cells – compounded by anaphase’s transient nature – and the inability to reconstitute mammalian spindles in vitro. Defining how mechanical forces flow through the anaphase spindle will be key to better understanding the physical principles driving its function.

The spindle midzone is a key structural module responsible for much of the mechanics of anaphase. In mammalian cells, antiparallel interpolar spindle microtubules are crosslinked by PRC1 (Subramanian *et al*., 2010) forming bundles with plus-end antiparallel overlaps in the center of the spindle (Euteneuer and McIntosh, 1980; Mastronarde *et al*., 1993; Mollinari *et al*., 2002; Conway *et al*., 2026). At anaphase, microtubules within these bundles slide against each other, actively generating outward force to drive chromosome segregation and spindle elongation (Vukušić *et al*., 2017; Yu *et al*., 2019; Chen *et al*., 2026). This sliding is thought to be driven by the kinesin-4 motor KIF4A and kinesin-5 motor Eg5 (Vukušić *et al*., 2021). Additionally, these bundles and associated overlap proteins provide passive frictional force to resist astral pulling forces and hypersegregation (Aist *et al*., 1993; Collins *et al*., 2014; Pamula *et al*., 2019), helping maintain the central spindle.

The presence of a force-generating structure in the spindle does not necessarily imply that it is responsible for chromosome segregation. Instead, contribution to chromosome movement at anaphase depends on how such structures are connected to one another and ultimately to chromosomes (Anjur-Dietrich *et al*., 2021). Recent work using laser ablation has probed the degree to which antiparallel anaphase midzone bundles are mechanically connected to each other, suggesting there are connections that reinforce the midzone while also mechanically isolating individual bundles from local perturbations (Carlini *et al*., 2022). However, it remains unclear how forces flow through the anaphase spindle over different directions, distances, and timescales. We also do not know the relative strength of different anaphase modules or their various connections, or even which connections can or cannot bear load. Ultimately, it remains unclear if these bundles exist as isolated units or if they function as a coordinated globally connected structural unit. Differentiating between these possibilities is essential to define mechanistic models of anaphase chromosome segregation and to understand what role, if any, these bundles play in helping resolve lagging chromosomes. Answering these questions requires the ability to directly apply local forces on these anaphase bundles with spatial and temporal control. Classic work has used microneedle manipulation to apply force to probe meiotic spindle mechanics in grasshopper spermatocytes (Nicklas *et al*., 1982) and frog extract systems (Shimamoto *et al*., 2011). Our lab has now brought this approach to mammalian mitotic cells (Long *et al*., 2020; Suresh *et al*., 2020) where we have probed the mechanics of the metaphase spindle, a steady-state structure. Doing so at anaphase remained more challenging since it requires fast manipulation due to the short lifetime and rapid structural transformation of the mammalian anaphase spindle.

Here, we establish the use of microneedle manipulation in mammalian anaphase spindles to probe its mechanical architecture. We show that we can robustly exert local, reproducible force on the anaphase spindle, and that we can apply force directly to midzone bundles. Our work demonstrates that midzone bundles are strongly connected to each other in the short spindle axis, transmitting force up to 4 µm away. Further, we find that applying transient force to a small fraction of bundles can stall and even reverse whole spindle elongation. This demonstrates the relative strength of both plus-end overlaps and midzone bundle minus-end connections to the spindle pole region, revealing the anaphase spindle’s structural priorities. This response is time-dependent and titratable, with slower forces applied over a longer time slowing down but not reversing spindle elongation. Finally, we demonstrate the microtubule bundler PRC1 is necessary for maintaining a strong coupling between both poles during anaphase, indicating limited mechanistic redundancy. Overall, this work shows that despite its transient and self-remodeling nature, the anaphase spindle remains a highly connected and robust structure across directions and timescales. We propose that such load-bearing connections could support coordinated segregation during anaphase, instead of each bundle acting as an isolated unit.

## Results

### Microneedle manipulation can exert local forces with spatiotemporal control on the mammalian anaphase spindle

Our lab recently adapted microneedle manipulation to mammalian mitotic cells (Long *et al*., 2020; Suresh *et al*., 2020), demonstrating that we can apply local loads to individual microtubule bundles with high spatial and temporal precision. We showed that we can do so without piercing the cell cortex or membrane, thus preserving cell health. Our efforts have so far focused on metaphase spindles whose long-lived structure made them an ideal candidate given that rapid needle movements can lead to membrane rupture. Here, we sought to bring microneedle manipulation to the mammalian anaphase spindle. We used PtK2 GFP-α-tubulin cells (Khodjakov *et al*., 2003) as their large, flat shape and low chromosome number make them ideal for spindle manipulations, and for pulling on individual microtubule bundles. We sought to directly apply force on midzone microtubule bundles to probe how the anaphase spindle transmits force in space and time. Across space, we aimed to probe the strength of midzone bundles’ overlaps, their connections to each other, and their connections to the rest of the spindle (Fig. 1 A). Across time, we aimed to probe how the anaphase spindle responds to short- vs long-lived forces by pulling the same distance over different timescales (Fig. 1 A). We used a fluorescently coated (BSA-Alexa 647) glass microneedle of 1 µm tip diameter bent to contact the cell at a 90° angle. We mounted it to a computer-controlled micromanipulator to achieve reproducible movements (Fig. 1 B). We synchronized cells with CDK1 inhibitor RO-3306 (Vassilev *et al*., 2006), followed by a washout, to enrich for mitotic cells. When a cell entered anaphase, determined visually by chromatid separation, we focused on that cell for manipulation, imaging using fluorescence spinning disk confocal microscopy every 5 s (Fig. 1 C).

**Figure 1.**
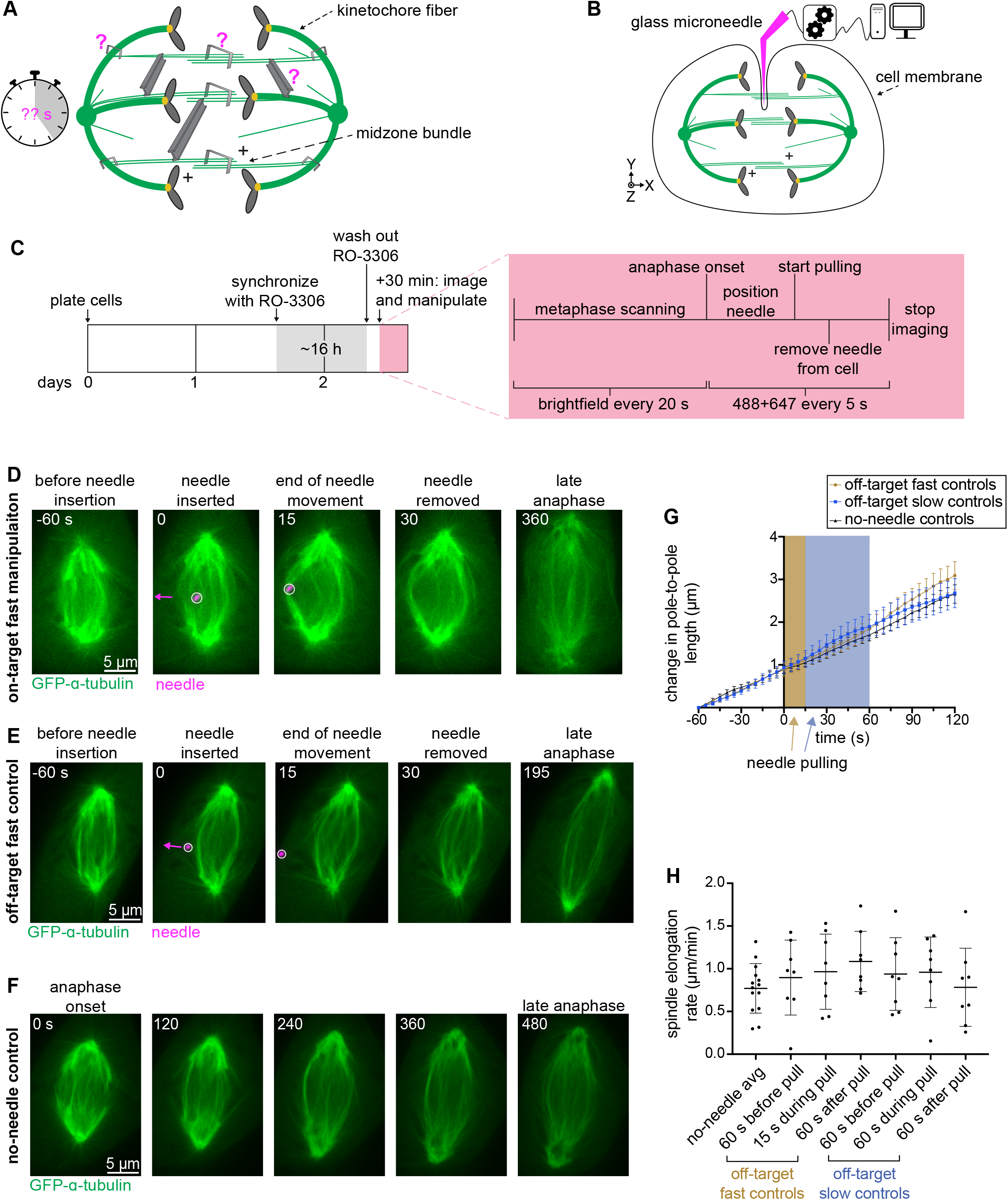
Microneedle manipulation can exert local forces with spatiotemporal control on the mammalian anaphase spindle. **(A)** Cartoon depicting anaphase spindle and midzone bundle connections that we aim to probe (magenta question mark) in this work: mechanics of antiparallel microtubule (green, with microtubule plus-ends labelled) overlaps (large staples), of midzone bundle to spindle connections (small staples), and of bundle-to-bundle connections (beams). We also aim to probe how these connections respond to forces of varying timescales (stopwatch). **(B)** Schematic illustration of computer-controlled (computer, with XYZ manipulator) microneedle (magenta) setup for midzone bundle manipulation assays. Microneedles do not rupture the cell membrane in our assay. **(C)** Schematic diagram of microneedle experiments. Cells were synchronized in G2 with CDK1 inhibitor RO-3306 for ∼16 h before imaging on confocal microscope (detailed in red box). **(D-E)** Representative timelapse confocal images of a PtK2 anaphase spindle (GFP-α-tubulin, green) during a 12.8s “on-target” fast manipulation (inside the spindle body, D) and “off-target” fast control (outside the spindle body, E) with microneedle (BSA Alexa Fluor 647, magenta, arrow shows direction of needle movement) displayed on images. Time = 0 s defined as frame before computer-controlled needle movement begins. **(F)** Representative timelapse confocal images of a PtK2 no-needle control anaphase spindle (GFP-α-tubulin, green). **(G)** Spindle length over time for no-needle (black triangle, n = 14), off-target fast (brown circles, n = 8), and off-target slow (blue squares, n = 8) controls. Shaded areas represent period of needle movement (brown = fast, blue = slow). Initial spindle length taken from first frame of imaging for no-needle controls and 60 s prior to pulling for off-target controls. Mean ± SEM. **(H)** Spindle elongation rate in no-needle controls (n = 14) averaged over first five minutes of anaphase, fast (n = 8) and slow (n = 8) off-target controls before, during, and after pulls. No significant difference between no-needle control and any other group of off-target pulls (Mann-Whitney test). Mean ± SD.

Given that the window from chromatid separation to the start of cytokinetic furrowing lasts about ten minutes in these cells, the first challenge was to position the microneedle inside the spindle fast enough to perform a computer-controlled pull before the start of cytokinesis. We found that pulls on a small subset of outer midzone bundles, and sometimes one bundle, were feasible during spindle elongation (Fig. 1 D and Video 1). In all pulls, we sought to position the needle near the spindle midzone next to an outer bundle, and to pull orthogonal to the spindle’s long axis. We established two pulling speeds: a “slow” speed of ∼6 µm over 60 s (4.6 ± 1.0 µm/min ) and a “fast” speed of ∼5 µm over 12.8 s (20.0 ± 1.5 µm/min) to probe the time-dependence of spindle responses to force. Our slow speed is near the lower limit allowed by our system while maintaining smooth movement and within an order of magnitude of spindle elongation rates (Fig. 1 H). Our fast speed applies a more transient disruption, moving the same distance over a shorter time span. Importantly, after removing the needle from the cell post-pull, most spindles continued through anaphase. When we imaged for many minutes after manipulation, cells completed cytokinesis. To control for adverse or indirect effects of microneedle perturbations on anaphase spindle function, we performed the same manipulations just outside the spindle (“off-target”) with the same parameters as our experimental perturbations (Fig. 1 E and Video 2). Prior to all analyses, movies were registered using tubulin signal to control for variability in whole cell or whole spindle translations. Spindle elongation speed was indistinguishable between cells with off-target manipulations and unmanipulated cells (Fig. 1 F-H and Video 3). These findings indicate that the structural responses we observe when exerting force inside the spindle result from force on spindle microtubules specifically, not from indirect effects of manipulation. Thus, we can use microneedle manipulation to exert local forces with spatial and temporal control on mammalian anaphase midzone bundles.

### Anaphase midzone bundles can transmit lateral force to each other over microns and tens of seconds

Pioneering work by Bruce Nicklas used microneedles to probe the connections of <u>k</u>inetochore-fibers (k-fiber) during anaphase in grasshopper spermatocytes, finding they are laterally coupled (Nicklas *et al*., 1982). Recent work in mammalian cells suggests that anaphase midzone bundles are mechanically isolated from their neighbors, even while being directly connected via crosslinking microtubules (Carlini *et al*., 2022). However, it remains unclear whether anaphase midzone bundles can transmit force between each other along the spindle’s short axis, and if so, over what timescales. Defining this timescale will be key to understanding how the anaphase spindle can remodel itself while maintaining its overall stability (Shimamoto *et al*., 2011).

To probe lateral force transmission between bundles, we performed microneedle manipulations of an outer bundle, measured how much other bundles moved as a result, and compared to “off-target” control manipulations (Fig. 2 A). For fast pulls, we moved the microneedle 5.1 ± 1.0 µm over 12.8 ± 1.5 s (23.8 ± 2.9 µm/min) (Fig. 2 B and Video 4). We calculated bundle displacement in the axis of needle movement based on the shortest pre- and post-manipulation distance to the pole-to-pole axis, normalizing to the needle’s displacement in that cell. With fast pulls, we observed strong correlated displacement of neighbor bundles in the same direction as the manipulated bundle (Fig. 2 C). We fit the displacement vs bundle distance curve to a single exponential, valid for any structure connecting bundles with a constant start and end probability over space, finding that displacement reached zero at a starting distance of 4.2 µm from the manipulated bundle based on the line of best fit. Similarly, when binning neighbor bundles by their distance to the manipulated bundle prior to pulling, we found significant coupling for bundles 0-4 µm away from the manipulated bundle (Fig. 2 D). This group of bundles moved significantly away from the pole-to-pole axis in the direction of needle movement when compared to off-target control. In contrast, bundles which started 4-8 µm away from the manipulated bundle moved significantly away from the pole-to-pole axis in the opposite direction (Fig. 2 D). This suggested to us that manipulation may result in a compressive force causing bundles to buckle outward. If these movements were only a result of compressive force, we would expect similar levels of outward buckling for bundles in each spindle half. However, when we binned bundles based on which spindle half they started on, we saw significantly more outward movement on the same side as the needle (Fig. 2 E). Therefore, we conclude that while buckling likely plays a role in bundle movement, some movement is caused by connections to the manipulated bundle.

**Figure 2.**
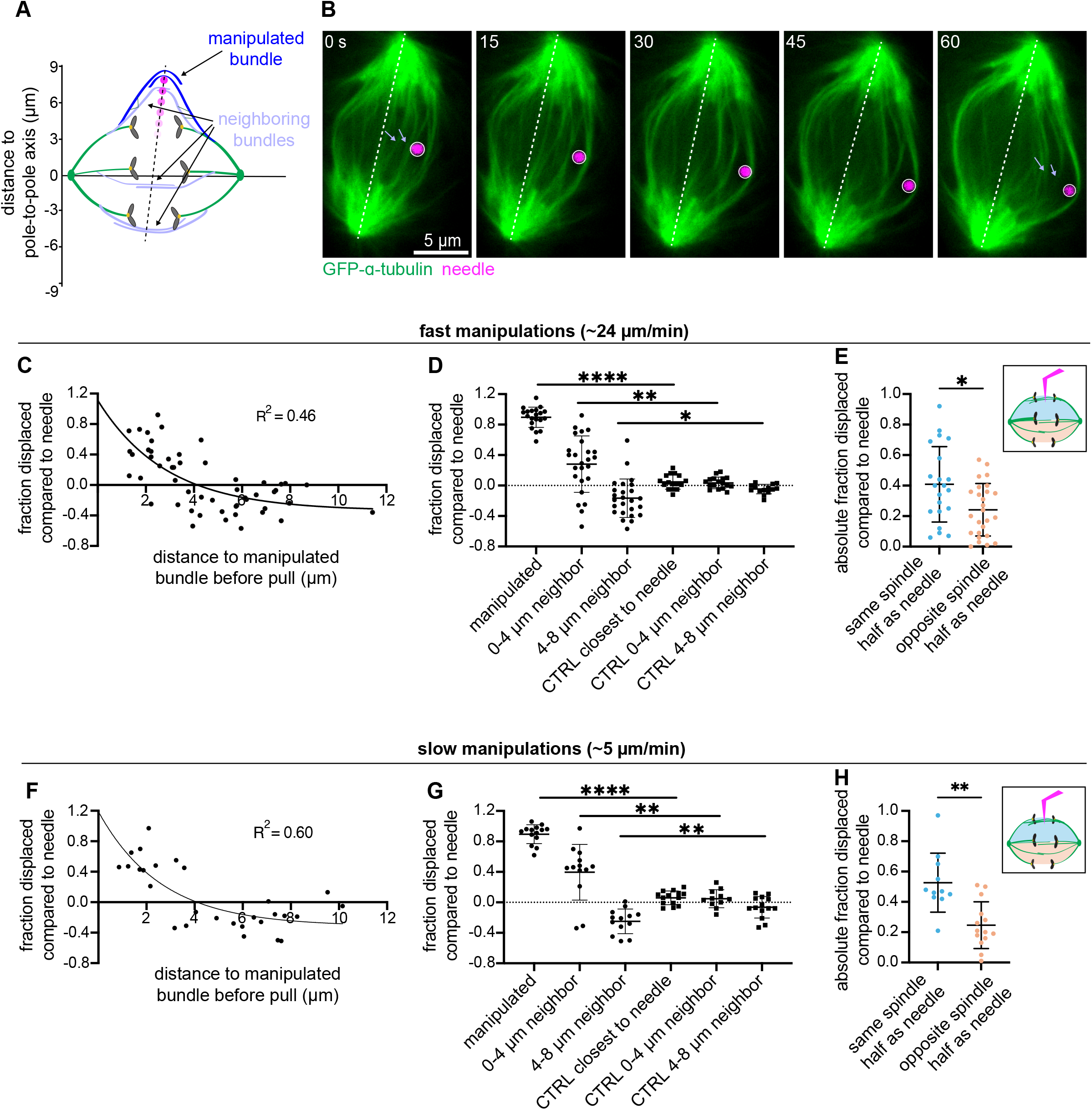
Anaphase midzone bundles can transmit lateral force to each other over microns and tens of seconds. **(A)** Schematic illustration of experiment design. We use the microneedle (magenta circle) to pull on a midzone bundle (dark blue, top) and measure the resulting displacement of other midzone bundles (light blue) along the spindle’s short axis. Measurement points on bundles were determined based on axis of needle movement (black dashed line). **(B)** Representative timelapse confocal images of a PtK2 anaphase spindle (GFP-α-tubulin, green) during a 60 s slow manipulation, with microneedle (BSA Alexa Fluor 647, magenta) displayed on images. Light blue arrows point to measured positions of two neighbor bundles pre- and post-manipulation. **(C)** Fast manipulation (5.1 ± 1.0 µm over 12.8 ± 1.5 s at 23.8 ± 2.9 µm/min) neighbor midzone bundle displacement normalized to needle displacement (fraction displaced compared to needle) versus distance from the manipulated bundle prior to pulling (n = 52 from 20 cells). Solid black line is the single exponential decay nonlinear regression fit to the data (r^2^ = 0.46). All measurements taken after registering tubulin channel to account for whole spindle and cell translation. **(D)** Fraction displaced compared to needle for fast manipulated bundles (n = 20 from 20 cells), and neighbor bundles binned by distance to manipulated bundle prior to pulling: 0-4 µm neighbors (n = 25 from 18 cells), 4-8 µm neighbors (n = 25 from 19 cells). Controls include bundle closest to needle (n = 16 from 16 cells) and neighbors binned by distance to closest bundle prior to pulling: CTRL 0-4 µm neighbors (n = 17 from 12 cells), CTRL 4-8 µm neighbors (n = 14 from 12 cells). Manipulated vs CTRL closest to needle (****p < 0.0001, Mann-Whitney test), 0-4 µm neighbors vs CTRL 0-4 µm neighbors (**p = 0.002, Mann-Whitney test), 4-8 µm neighbors vs CTRL 4-8 µm neighbors (*p = 0.01, Mann-Whitney test). Mean ± SD. **(E)** Fast manipulation absolute fraction displaced compared to needle for neighboring bundles binned by spindle half (same as needle n = 21, opposite from needle n = 26) prior to pulling, plotted based on displacement from pole-to-pole axis during pull (*p = 0.019, Mann-Whitney test). Mean ± SD. **(F)** Slow manipulation (4.6 ± 0.9 µm, over 60.7 ± 7.3 s at 4.9 ± 0.8 µm/min) neighbor midzone bundle displacement normalized to needle displacement (fraction displaced compared to needle) versus distance from the manipulated bundle prior to pulling (n = 28 from 14 cells). Solid black line is the single exponential decay nonlinear regression fit to the data (r^2^ = 0.60). **(G)** Fraction displaced compared to needle for slow manipulated bundles (n = 14 bundles from 14 cells), and neighbors binned by distance to manipulated bundle prior to pulling: 0-4 µm neighbors (n = 13 from 13 cells), 4-8 µm neighbors (n = 13 from 13 cells). Controls include bundle closest to needle (n = 14 from 14 cells) and neighbors binned by distance to closest bundle prior to pulling: CTRL 0-4 µm neighbors (n = 11 from 11 cells), CTRL 4-8 µm neighbors (n = 14 from 13 cells). Manipulated vs CTRL closest to needle (****p < 0.0001, Mann-Whitney test), 0-4 µm neighbors vs CTRL 0-4 µm neighbors (**p = 0.0039, Mann-Whitney test), 4-8 µm neighbors vs CTRL 4-8 µm neighbors (**p = 0.0068, Mann-Whitney test). Mean ± SD. **(H)** Slow manipulation absolute fraction displaced compared to needle for neighboring bundles binned by spindle half (same as needle n = 11, opposite from needle n = 14) prior to pulling, plotted based on displacement from pole-pole axis during pull (***p = 0.0013, Mann-Whitney test). Mean ± SD.

To probe the timescales over which these connections remain, we then performed slower manipulations pulling a similar distance as fast pulls, 4.6 ± 0.9 µm, over 60.7 ± 7.3 s (4.9 ± 0.8 µm/min), near the lower speed limit of smooth needle movement achievable with this manipulator. Here too, we observed strong lateral force transmission, with bundle displacement reaching zero at a neighbor starting distance of 4.1 µm (Fig. 2 F). We also find that the transformed slopes of the regression curves for fast and slow pulls are statistically similar (Fig. S1 C). In these slow pulls, there was again significant concordant displacement of neighboring bundles that started 0-4 µm away from the manipulated bundles (Fig. 2 G). For these slow pulls, the lateral movements of neighbors were significantly different between each spindle half (Fig. 2 H). This demonstrates that movements on the needle-proximal side of the spindle are due to connections to the manipulated bundle and not simply due to compressive buckling. Together, these findings demonstrate that anaphase microtubule bundles are strongly connected and can bear load laterally, transmitting force across nearly half the spindle for up to at least one minute.

### Transient force on midzone bundles reverts whole anaphase spindle elongation, demonstrating strong longitudinal coupling of anaphase components on short timescales

We next set out to define force transmission along anaphase midzone bundles in the spindle’s long axis – the spindle’s natural force generation axis (Fig. 3 A). We reasoned that our orthogonal force could be dissipated locally by i) bundles elongating through sliding, ii) bundles rupturing, as we observed over minutes at metaphase (Long *et al*., 2020; Rux *et al*., 2026), or iii) bundle ends detaching from their spindle anchor points. Alternatively, if midzone bundles don’t significantly elongate, rupture, or detach from their anchor points under force, iv) the spindle could dissipate applied force globally by changing its shape, e.g. through inward movement of spindle poles (Fig. 3 B).

**Figure 3.**
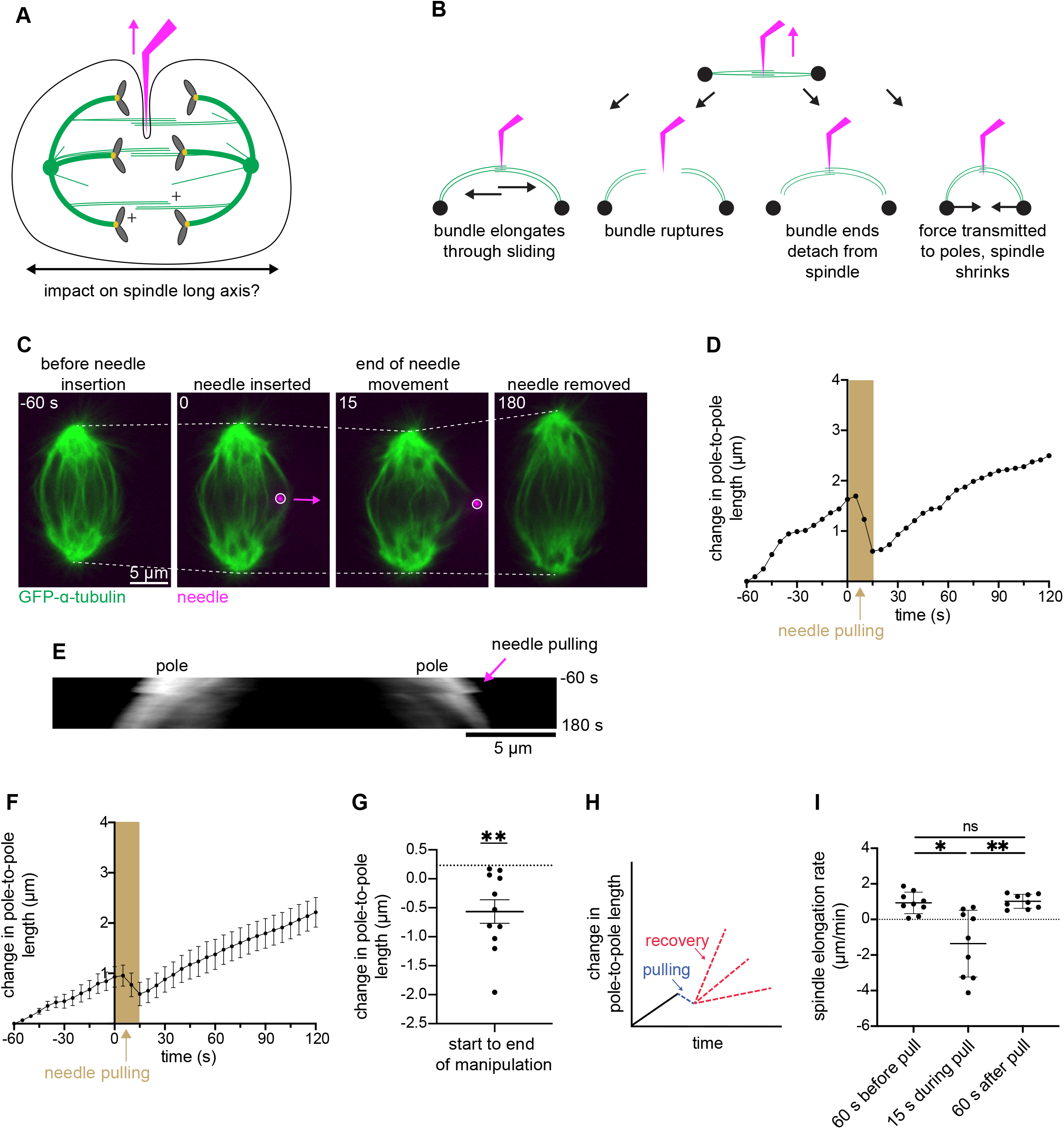
Transient force on midzone bundles reverts whole anaphase spindle elongation, demonstrating strong longitudinal coupling of anaphase components on short timescales. **(A)** Schematic illustration showing direction (magenta arrow) of microneedle (magenta) manipulation and direction of force propagation assessed in this figure (black arrow). **(B)** Models of midzone bundle response to external force: bundle elongates through sliding; bundle ruptures; bundle ends detach from spindle; force is transmitted to poles and spindle shrinks. **(C)** Representative timelapse confocal images of a PtK2 anaphase spindle (GFP-α-tubulin, green) during a 12.8 s fast manipulation (4.7 ± 0.5 µm at 22.2 ± 2.5 µm/min), with microneedle (BSA Alexa Fluor 647, magenta) displayed on images, arrow shows direction of needle movement. Dashed lines show spindle length used in measurements. **(D)** Spindle length versus time for the cell in (C). Time = 0 s is the frame before needle pulling and the shaded area represents duration of needle movement. Initial spindle length taken 60 s prior to needle pulling. **(E)** Kymograph of spindle from (C) and (D), (GFP-α-tubulin, grey), with needle pulling (pink arrow) occurring at time = 0 s, followed by spindle shortening. **(F)** Spindle length change versus time for all fast manipulations (n = 9). All cells aligned to time = 0 s. Change in length compared to spindle length 60 s prior to pull. Mean ± SEM. **(G)** Change in spindle length from start to end of manipulation for fast manipulations (n = 11). Mean is compared to expected change based on average pre-pull elongation rate (dotted line) (**p = 0.001 Wilcoxon signed rank test). Mean ± SEM **(H)** Schematic showing possible outcomes after spindle shrinkage due to needle force application (blue): the spindle could either slow down its elongation due to damage (red bottom), continue at the same elongation rate as before manipulation (red middle), or elongate faster after storing force generation potential (red top). **(I)** Spindle elongation rate in fast manipulations (n = 9) before, during, and after pulls. No significant difference between pre- and post-pull elongation rates (p = 0.65, Wilcoxon test). During pull is different than pre- and post-pull elongation rates (*p = 0.01, **p = 0.008, Wilcoxon matched-pair signed rank test). Mean ± SD.

We pulled on midzone bundles 4.7 ± 0.5 µm over 12.8 s (22.2 ± 2.5 µm/min), a much shorter time span than the minutes scale of anaphase spindle elongation. Though imaging limitations make it difficult to determine in all pulls the extent to which force may be dissipated via bundle sliding, rupture, or detachment, we found that sufficient force was transmitted along midzone bundles to stall and even reverse spindle elongation (Fig. 3 C-E and Video 5), in most spindles (7/11) we pulled on (Fig. 3 F and G). This demonstrates continuous longitudinal force transmission from pole to pole during anaphase. Additionally, our findings reveal the relative strength of these midzone bundles and their connection points: their overlaps can resist large outward force sufficient to shorten the whole spindle, while the entire bundle remains robustly attached to the rest of the spindle. These findings reveal a structural priority of the anaphase spindle, where transient reversal of global spindle elongation is favored over disruption of the local central spindle architecture.

Next, we asked how the spindle responds to this force application and spindle shrinkage after needle and force removal (Fig. 3 H). Anaphase spindles could produce more force in response to obstruction (Nicklas, 1983), increasing elongation rate to “catch up” to their expected length, they could maintain the same pre-pull elongation rate unaffected by force application and spindle shrinkage, or the elongation rate could decrease if force generating structures had been destroyed and must remodel before normal elongation can continue. To distinguish between these scenarios, we compared the spindle elongation rate before and after manipulation and found that pre- and post-pull elongation rates were not significantly different (Fig. 3 I). This suggests that the spindle did not respond to this disruption by upregulating elongation force and indicates that the spindle’s force generation ability was not impacted by external force application. Interestingly, the return to the pre-pull elongation rate appears almost instantaneous (Fig. 3 F). This indicates that the force production unit of anaphase spindle elongation is robust to, and does not adapt to, these structural and mechanical disruptions. Together, these findings reveal the relative strength of midzone bundle overlaps and their spindle connections and demonstrate the architectural priority of the anaphase spindle to preserve midzone overlap structure and the robustness of spindle elongation to structural and mechanical fluctuations.

### Sustained force on midzone bundles transiently slows anaphase spindle elongation

To achieve its function, the anaphase spindle must maintain its structure over shorter timescales and remodel its structure over longer ones. As such, its material properties may be time-dependent. We know that the *Xenopus* extract metaphase spindle’s response to external forces is viscoelastic and thereby dependent on the timescale of force application (Shimamoto *et al*., 2011). Similarly, in vitro work on minimal antiparallel microtubule systems has shown variable frictional responses to applied forces of differing speeds (Shimamoto *et al*., 2015; Gaska *et al*., 2020; Alfieri *et al*., 2021). Thus, we asked if, and how, the mammalian anaphase spindle responds differently to slower and more prolonged pulling forces.

We again applied force directly to outer midzone bundles, pulling 4.9 ± 0.8 µm over 60 s (4.9 ± 0.8 µm/min), 4-5 times slower than in the above fast pulls (Fig. 3). However, unlike in the fast pulls, we did not see spindle elongation reversal (Fig. 4 A-E and Video 6). Instead, spindle elongation rate only slowed during force application which led to a smaller spindle length post-pull than expected based on the pre-pull elongation rate (Fig. 4 F). As in response to fast pull (Fig. 3 I), we observed no significant change in spindle elongation rates comparing pre- and post-pull (Fig. 4 G). Interestingly, we observed four spindles (across two experimental days) with elongation rates that were faster post-pull than pre-pull, and faster than rates we observed in any other conditions (Fig. 4 G). While these observations hint at the possibility of an active response compensating for the “lost” elongation, we cannot make such a conclusion at this time. Thus, anaphase midzone bundle overlaps and their anchorage to the rest of the spindle are not only structurally robust but can persist in response to forces of variable speeds and timescales.

**Figure 4.**
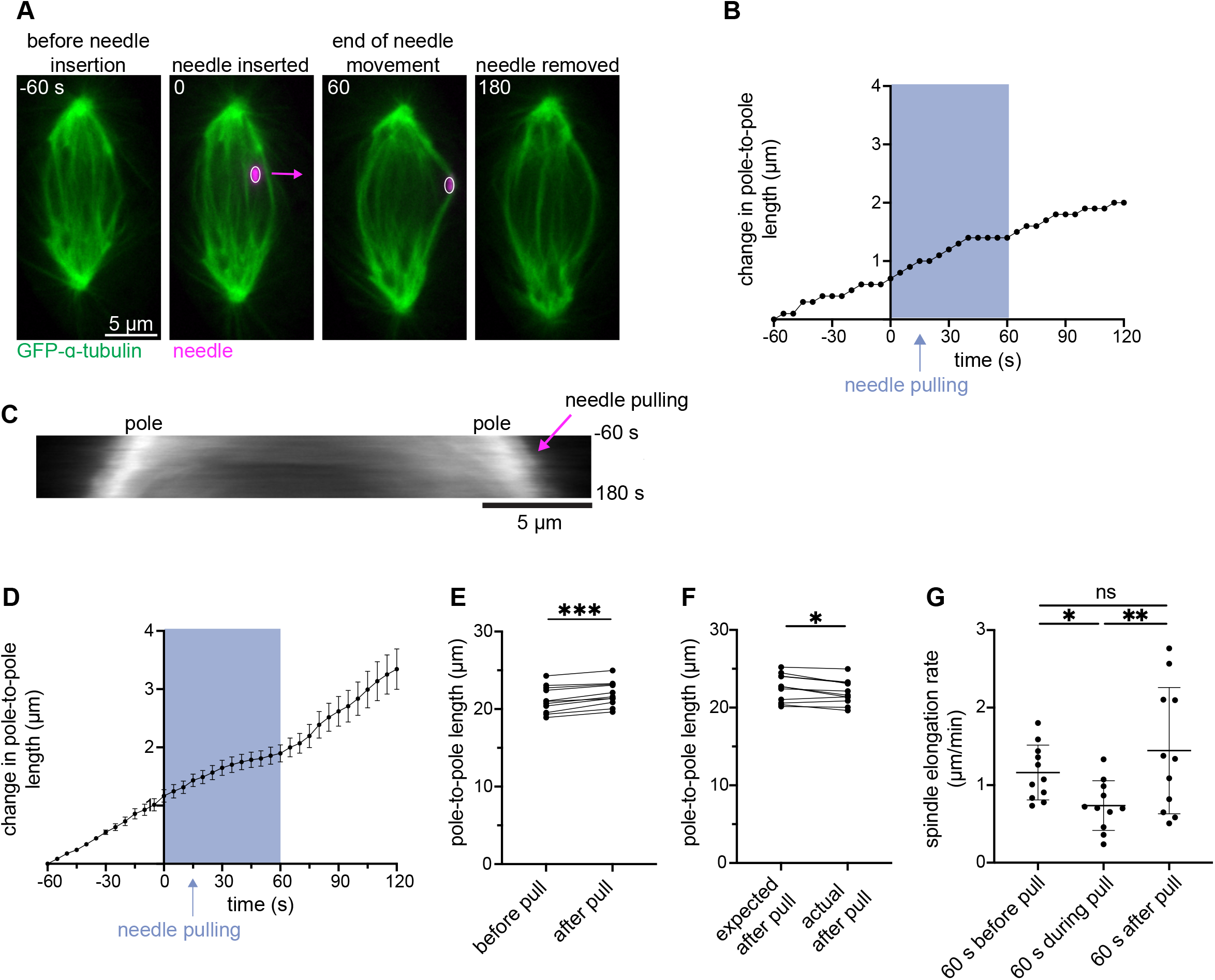
Sustained force on midzone bundles transiently slows anaphase spindle elongation. **(A)** Representative timelapse confocal images of a PtK2 anaphase spindle (GFP-α-tubulin, green) during a 60 s slow manipulation (4.9 ± 0.8 µm at 4.9 ± 0.8 µm/min), with microneedle (BSA Alexa Fluor 647, magenta) displayed on images, arrow shows direction of needle movement. **(B)** Spindle length versus time for cell in (A). Time = 0 s is the frame before needle pulling and the shaded area represents duration of needle movement. Initial spindle length taken 60 s prior to needle pulling. **(C)** Kymograph of spindle from (A), (GFP-α-tubulin, grey), with needle pulling (magenta arrow) occurring at time = 0 s. **(D)** Spindle length change versus time for all slow manipulations (n = 11). All cells aligned to time = 0 s. Shaded area represents duration of needle movement, initial spindle length taken 60 s prior to needle pulling. Mean ± SEM. **(E)** Absolute spindle length in frame before and after slow manipulation (n = 11, ***p = 0.001, Wilcoxon matched-pair signed rank test). **(F)** Expected spindle length (based on pre-pull elongation rate) and actual spindle length after slow manipulation (n = 11, *p = 0.02, Wilcoxon matched-pair signed rank test). **(G)** Spindle elongation rate in slow manipulations (n = 11) before, during, and after pulls. No significant difference between pre- and post-pull elongation rates (p = 0.70, Wilcoxon test). During pull is different than pre- and post-pull elongation rates (*p = 0.0186 and **p = 0.0049 respectively, Wilcoxon test). Mean ± SD.

### PRC1-mediated crosslinking maintains a force transmitting connection between spindle poles during anaphase

The metaphase spindle has been described as a unitary body, maintaining a closed force loop between poles to move chromosomes (Pereira and Maiato, 2012). Our manipulations of anaphase spindles demonstrate that even by mid-anaphase, spindle structures continue to maintain this strong mechanical connection between spindle poles (Fig. 3). We asked if this force transmission was specifically due to antiparallel midzone bundles of microtubules, or if it relied on redundant mechanisms, for example from a non-specific network of interconnected spindle microtubules (Mastronarde *et al*., 1993; Redemann *et al*., 2017; O’Toole *et al*., 2020). To target midzone bundles, we used siRNA to reduce expression of PRC1, a crosslinking protein important for the formation and maintenance of these bundles (Mollinari *et al*., 2002; Polak *et al*., 2017). We assayed knockdown via immunofluorescence and confirmed that PRC1 was significantly reduced (Fig. 5 A and B). Additionally, we could typically no longer clearly identify distinct midzone bundles (Fig. 5 C and Video 7). We performed manipulations similar to fast pulls (as in Fig. 3), pulling microneedles over 4.5 ± 0.8 µm for 12.8 s (21.0 ± 3.6 µm/min). We found that these spindles no longer detectably shrunk in response to manipulation force (Fig. 5 D, E, and F). While we do observe a small but significant reduction in spindle length compared to the expected length without manipulation (Fig. 5 G), the extent of this reduction was markedly smaller when compared to WT cells with PRC1 (Fig. 5 H). The remaining response may result from incomplete PRC1 knockdown (Fig. 5 B) or remaining microtubule crosslinking via Eg5 (Conway *et al*., 2026), for example. Further, we observed a small decrease in spindle elongation rate during manipulation, and interestingly after manipulation as well (Fig. 5 I). These findings demonstrate that until at least mid-anaphase, PRC1-mediated antiparallel midzone bundles are essential for maintaining the spindle as a closed unitary body, with force transmitted pole-to-pole, and indicates limited mechanical redundancy coupling both anaphase spindle halves.

**Figure 5.**
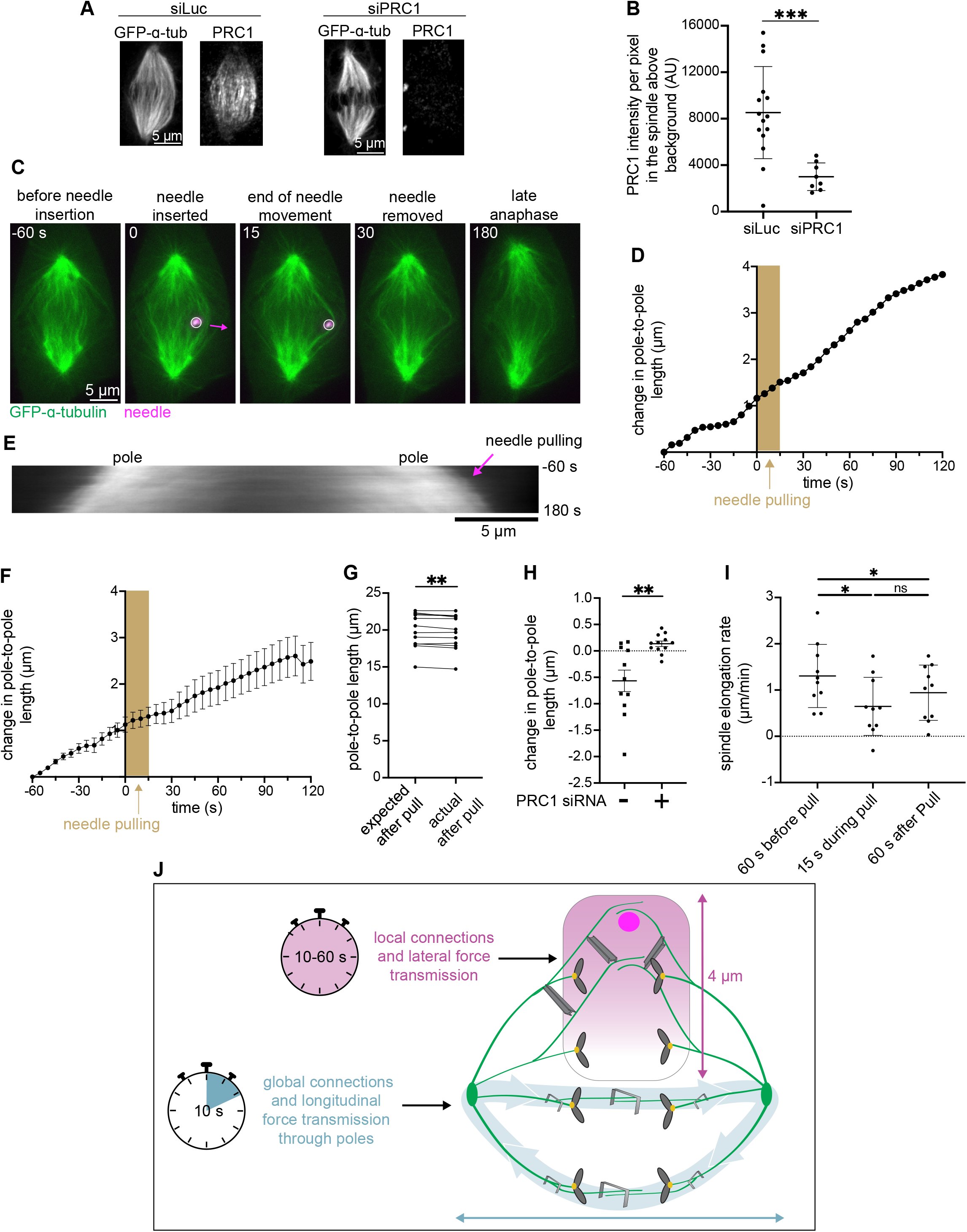
PRC1-mediated crosslinking maintains a force transmitting connection between spindle poles during anaphase. **(A)** Representative immunofluorescence confocal images of a PtK2 GFP-α-tubulin anaphase spindle treated with siRNA against luciferase or PRC1 for 72 h. **(B)** PRC1 signal quantification from immunofluorescence of anaphase cells treated with siLuc (n = 15) or siPRC1 (n = 8). PRC1 intensity is significantly different between groups (***p = 0.0008, Mann-Whitney test). Mean ± SD. **(C)** Representative timelapse confocal images of a PtK2 anaphase spindle (GFP-α-tubulin, green) treated with siRNA against PRC1 during a 12.8s fast manipulation (4.5 ± 0.8 µm at 21.0 ± 3.6 µm/min), with microneedle (BSA Alexa Fluor 647, magenta) displayed on images, arrow shows direction of needle movement. **(D)** Plot of spindle length versus time for cell in (C). Time = 0 s is the frame before needle movement and the shaded area represents duration of needle movement. Initial spindle length taken from 60 s prior to needle pulling. **(E)** Kymograph of spindle from (C) (GFP-α-tubulin, grey), with needle pulling (magenta arrow) occurring at time = 0 s. **(F)** Spindle length change versus time for all fast pulls (n = 10). All cells aligned to time = 0 s. Mean ± SEM. **(G)** Expected spindle length (based on pre-pull spindle elongation rate) and actual spindle length after 12.8s fast manipulation (n = 12, **p = 0.009 Wilcoxon test). **(H)** Change in spindle length from start to end of manipulation for fast pulls of cells with (n = 12) and without (n = 11) PRC1 siRNA (**p = 0.004, Mann-Whitney test). Mean ± SEM. **(I)** Spindle elongation rate in PRC1 siRNA fast pull cells (n = 10) before, during, and after pulls. Compared to the pre-pull elongation rate, both the during- and post-pull elongation rates are significantly different (*p = 0.02 and *p = 0.02 respectively, Wilcoxon test). No significant difference between during- and post-pull elongation rates (p = 0.2754, Wilcoxon test). Mean ± SD. **(J)** Model of the mammalian anaphase spindle’s mechanical architecture. Along the spindle’s short axis, force is transmitted between midzone bundles locally up to 4 µm away (magenta gradient and connecting beams). The strength of these connections drops off with distance and these connections can bear load up to at least one minute (magenta stopwatch). Along the spindle’s long axis, force is communicated globally from pole to pole (blue arrows). Force transmission requires strong overlaps and bundle connections to the rest of the spindle (grey staples) and is strongest over short timescales (blue stopwatch). We propose that these strong load-bearing connections across directions help ensure robust and coordinated separation of genetic material while allowing the spindle to remodel over long timescales.

## Discussion

The anaphase spindle must drastically deform, constantly generating and responding to force while still maintaining the structures necessary to complete cell division. Here, we use microneedle manipulation to probe the mechanical architecture of the anaphase spindle across space and time. Inspired by Nicklas’s classic work (Nicklas and Koch, 1969; Nicklas *et al*., 1982), we show that microneedle manipulation of anaphase spindles in live mammalian cells is possible during spindle elongation, does not significantly impact completion of anaphase, and induces direct structural responses (Fig. 1). Further, we demonstrate that anaphase midzone bundles are locally connected along the spindle’s short axis, with both transient (∼10 s) and prolonged (∼1 min) forces transmitting to bundles up to 4 µm away (Fig. 2). Midzone bundle overlaps and their connections to the rest of the spindle are robust, resisting excess local sliding, breakage, or detachment under microneedle manipulation. Instead, force is transmitted globally, from one pole to the other, stalling or even reversing spindle elongation in a time-dependent manner (Fig. 3 and 4), despite microtubules within these bundles not reaching the poles themselves (Mastronarde *et al*., 1993). Finally, we show that PRC1-mediated crosslinking is essential for maintaining this strong, long-range connection during anaphase (Fig. 5). Overall, these findings demonstrate that even as the anaphase spindle drastically deforms, it remains a mechanically connected structure both locally and globally across timescales and directions (Fig. 5 J). We propose that these strong load-bearing connections help ensure robust and coordinated separation of genetic material.

Whether antiparallel bundles in the mammalian spindle function as isolated units or as part of a single coordinated structure is not known and has clear implications for spindle function. Previous live imaging showed that PRC1-labeled bundles at metaphase have correlated movements with a single pair of sister kinetochores, suggesting these bundles act as individual units (Polak *et al*., 2017). It has also been proposed that these bundles remain mechanically isolated during anaphase (Carlini *et al*., 2022). Our findings here demonstrate that these PRC1-mediated bundles, at least at anaphase, are connected laterally along their length, with their connections bearing load over multiple timescales (Fig. 2). Also pointing to the anaphase spindle acting as a coordinated structure, force application to a small subset of bundles is sufficient to stall or reverse spindle elongation (Fig. 3 and 4). Prior modeling work led to the proposal that lateral connections between k-fibers could equalize forces on kinetochores at metaphase, ensuring coordinated chromosome segregation at anaphase (Matos *et al*., 2009). Similar force coordination between antiparallel bundles could support evenly distributed outward pushing forces that drive anaphase spindle elongation and chromosome to pole movement. Force coordination may also help bundle overlap regions stay aligned with one another, forming a tightly defined overlap zone and ensuring focused cytokinetic signaling. Future work will be needed to determine the molecular nature of these lateral connections and how their properties change as bundles become more stable and closer together as anaphase progresses (do Rosário *et al*., 2023). Finally, it will be exciting to test if these connections can support anaphase error correction mechanisms, for example with neighboring bundles transmitting force to each other to aid in the proper segregation of lagging chromosomes.

Whether the anaphase spindle adapts a “set-and-forget” trajectory or whether it adapts its trajectory to the force it experiences remains an open question. In the former model, spindle architecture built during metaphase and spindle assembly checkpoint satisfaction predetermine the motions of anaphase. In the latter model, the spindle adapts its structure and force generation potential to its force history. Favoring a “set-and-forget” model, we find that following needle removal, spindle elongation does not speed up to compensate for lost elongation (Fig. 3 I). Consistent with this, persistent lagging chromosomes slow down spindle elongation (Cimini *et al*., 2004). Mechanical isolation of midzone bundles may protect the spindle from one lagging chromosome disrupting them all (Carlini *et al*., 2022). Yet, hints of more complicated force-responsive mechanisms exist. When physically challenged, anaphase spindles in some systems appear to produce orders of magnitude more force than is normally required to separate chromatids (Nicklas, 1983). In vitro, overlap proteins can generate differential friction modes depending on force trajectories (Shimamoto *et al*., 2015; Gaska *et al*., 2020; Alfieri *et al*., 2021) and yet-to-be-tested feedback models propose how an adaptive midzone could stabilize overlaps under force (Hannabuss *et al*., 2019). Indeed, we observe a velocity-dependent response where preserving central spindle architecture is prioritized over transient reversal of global spindle elongation on short time scales (Fig. 3), while the spindle remodels more permissively under sustained loads (Fig. 4). Such a response could help the spindle remain robust in the face of transient forces, while elongating undisrupted around long-lived disruptions such as a persistent lagging chromosome. Microneedle manipulation could in principle help resolve these models, for example, by testing whether the changes to overlap protein dynamics during anaphase (Asthana *et al*., 2021) is purely driven by biochemistry or is force-responsive, and by directly testing an overlap-maintaining force feedback (Hannabuss *et al*., 2019).

Ultimately, anaphase stands as the last chance for chromosome segregation errors to be resolved or “locked in”. As such, probing the physical mechanisms driving anaphase spindle function is vital for understanding how failures arise. For example, the crosslinker PRC1, which is critical to maintaining a load-bearing connection between spindle poles during anaphase (Fig. 5), is highly dysregulated across cancer cell-types (Carter *et al*., 2006) and its overexpression correlates with increased proliferation, aneuploidy, metastasis, and drug resistance (Chen *et al*., 2016; Zhan *et al*., 2017; Li *et al*., 2018; Bu *et al*., 2020; Xu *et al*., 2020). Better understanding how altered PRC1 expression impacts local and global anaphase mechanics to facilitate tumor evolution or adaptively protect chromosomally unstable cancer cells will be an important future step.

## Acknowledgements

We thank Alexey Khodjakov for PtK2 GFP-α-tubulin cells. We are grateful to Wallace Marshall and Orion Weiner for helpful discussions, Caleb Rux for technical help, and members of the Dumont Lab for discussions and critical reading of the manuscript. This work was supported by NIH/NCI Predoctoral Fellowship F31CA291047 (ZMB), AHA Predoctoral Fellowship A143166 (ZMB), NIH R35GM136420 (SD), NSF 1548297 (SD) and the Chan Zuckerberg Biohub Investigator Program.

## Author Contributions

Conceptualization, Z.M-B., S.v.W., C.G., S.D.; methodology, Z.M-B.; validation, Z.M-B.; formal analysis, Z.M-B., S.v.W.; data curation, Z.M-B., S.v.W.; writing – original draft, Z.M-B.; writing - review and editing, Z.M-B., S.v.W., C.G., N.H.C., S.D.; visualization, Z.M-B., S.v.W.; supervision, S.D.; funding acquisition, Z.M-B. and S.D.

## Materials and methods

### Cell culture

All work herein was performed using PtK2 GFP-α-tubulin cells (gift from Alexey Khodjakov). PtK2 GFP-α-tubulin cells were cultured in MEM (11095-080; Thermo Fisher, Waltham, MA) supplemented with 1 mM sodium pyruvate (11360-070; Thermo Fisher), 1% non-essential amino acids (11140-050; Thermo Fisher), 1% penicillin/streptomycin (30-002-CI, Corning, Corning, NY) and 10% heat-inactivated fetal bovine serum (110438-026, Gibco, Waltham, MA). Cells were maintained at 37 °C and 5% CO_2_. The cell line tested negative for mycoplasma.

### Drug treatments

To enrich for cells in anaphase, we arrested cells before mitotic entry with the CDK1 inhibitor RO-3306 (Sigma-Aldrich, St. Louis, MO) at 4 µM for 16 h. RO-3306 was washed out 30 min prior to live imaging.

### Microscopy

Cells were imaged using a spinning disk confocal (CSU-X1; Yokogawa, Tokyo, Japan) inverted microscope (Eclipse Ti-E; Nikon instruments, Melville, NY) with a 100x 1.45 Ph3 oil objective through a 1.5X lens with the following components: head dichroic Semrock Di01-T405/488/568/647, 488 nm (150 mW) and 642 nm (100 mW) diode lasers (for tubulin and microneedle respectively), emission filters ET 525/50M and ET690/50M (Chroma Technology, Bellows Falls, VT), and a Zyla 4.2 camera (Andor Technology, Belfast, Northern Ireland), yielding 58 nm/pixel at bin = 1, or a Hamamatsu Quest2 qCMOS camera, yielding 46 nm/pixel at bin = 1. The microscope was operated by µManager (ver. 2.0.0. (Edelstein *et al*., 2014)).

### siRNA (Fig. 5)

For depletion of PRC1, cells were transfected with siRNA (5’-GGACTGAGGUUGUCAAGAA-3’) for PRC1 using Oligofectamine (Thermo Fisher, Waltham, MA) as previously described (Udy et al., 2015). Cells were imaged 72 hr after siRNA treatment. Knockdown was validated by counting binucleated cells and immunofluorescence analyses (Fig. 5 A and B). For live imaging, we confirmed PRC1 knockdown in the particular coverslips used by verifying at low magnification the enrichment of binucleated cells (from 2% (n = 200 cells) in luciferase control siRNA cells to 20.5% (n = 200) in PRC1 siRNA cells), a previously characterized consequence of PRC1 knockdown (Mollinari et al., 2002; Udy et al., 2015).

### Immunofluorescence to confirm PRC1 RNAi (Fig. 5 A and B)

We further confirmed PRC1 knockdown via immunofluorescence. Following siRNA treatment in PtK2 GFP-α-tubulin cells, cells were fixed with 95% methanol + 5 mM EGTA at -20°C for 5 min, washed with TBS-T (0.1% Tween20 in TBS), and blocked with 2% BSA in TBS-T for 1 h. Primary and secondary antibodies were diluted in TBS-T+2% BSA and incubated with cells for 1 h (primary) and for 45 min (secondary) at room temperature. DNA was labeled with Hoescht 33342 (Sigma, St. Louis, MO) before cells were mounted in ProLongGold Antifade (P36934; Thermo Fisher, Waltham, MA). Antibodies: rabbit anti-PRC1 H70 (1:100, Santa Cruz Biotechnology, Santa Cruz, CA), ChromoTek GFP-Booster Alexa Fluor 488 (1:200, Proteintech, Rosemont, IL), anti-rabbit secondary antibodies (1:500) conjugated to Alexa Fluor 647 (A21244; Life Technologies, San Francisco, CA).

Coverslips were imaged using the Nikon microscope described above. Z-stack images were acquired to include the entire spindle, and measurements were performed on sum-intensity projections. The average per pixel PRC1 intensity in the spindle measured above that in the cytoplasm was 8194 ± 1025 (AU) (SEM, n = 15) in luciferase control siRNA cells and 2704 ± 420 (AU) (SEM, n = 8) in PRC1 siRNA cells (67% knock-down, Fig. 5 B).

### Live imaging

For live imaging, cells were plated on glass-bottom 35 mm dishes coated with poly-D-lysine (P35GC-1.5-20-C; MatTek Corporation, Ashland, MA). Cells were imaged ∼48 h after plating on a stage-top incubation chamber (Tokai Hit, Fujinomiya-shi, Japan; or OkoLab, Pozzuoli, Italy) with the lid off to allow for microneedle manipulation. Cells were imaged at 30 °C and in CO_2_-independent media (18045-088; Gibco, Waltham, MA) + 10% heat-inactivated fetal bovine serum (10438-026, Gibco, Waltham, MA). Before selecting a live cell for imaging, we scanned multiple positions of metaphase cells using brightfield (300ms exposure every 20s) until one of them went into anaphase, or we selected cells in approximately the first 2 min of anaphase by morphological assessment of sister chromatid separation (Fig. 1 C). We then imaged cells with the 488 nm and 640 nm excitation lasers and brightfield every 5 s during the timeframe of manipulation (Fig. 1 C). All image analysis was performed during this second imaging window. For some cells analyzed in Fig. 2, this second imaging window included 5 z-stacks (0.5 µm step size) to improve throughput of neighbor bundle identification. All other imaging was performed in a single z-plane.

### Microneedle manipulation

Manipulations were performed as previously described (Suresh et al., 2020) unless stated otherwise.

- **Making microneedles:** Glass capillaries with an inner and outer diameter of 1 mm and 0.58 mm respectively (1B100-4 or 1B100F-4, World Precision Instruments) were used to create microneedles. A micropipette puller (P-87, Sutter Instruments, Novato, CA) was used to create uniform glass microneedles. Microneedles were bent ∼1.5 mm away from their tip to a 45° angle using a microforge (Narishige International, Amityville, NY), so as to have them approach the coverslip at a 90 ° angle (the microneedle holder was 45° from the coverslip). Our needles were dyed using a BSA Alexa Fluor 647 conjugate (BSA-Alexa-647; A-34785, Invitrogen).
- **Choosing cells:** We marked metaphase cells if they were flat and had bipolar spindles with both poles in the same focal plane. We tracked cells in brightfield until we observed a cell go into anaphase. Anaphase cells were chosen for manipulation if they had maintained a flat shape with an in-plane bipolar spindle.
- **Contacting the cell with microneedles:** To aid in placing the needle, the needle was left positioned above the coverslip at the center of the camera field of view (determined using µManager). Cells chosen for manipulation were positioned at this center point before starting the 5 s/frame imaging routine. Once an outer midzone bundle was identified, we used the needle fluorescence to guide its position. This positioning was done manually using the 3-axis-knob controller (ROE-200, Sutter Instruments, Novato, CA). Through this iterative process, we positioned the microneedle such that it was inside the spindle, right next to an outer midzone bundle.
- **Moving microneedles:** Once the microneedle was positioned next to an outer midzone bundle, it was moved in a direction roughly perpendicular to the spindle’s pole-to-pole axis. These manipulations were performed with computer control that took the following inputs: angle of movement, duration of movement, and distance. The needle was then lifted out of the cells following manipulation either manually using the 3-axis-knob-controller or automatically via a Python script. Controls were performed by placing the needle just outside the spindle, next to the outer midzone bundle and performing the manipulation in the same manner as experimental manipulations as described above. Fast manipulation input parameters were set to move the needle and absolute distance of ∼5 µm over 10-15 s and slow parameters were set to move the needle ∼6 µm over 50-70 s with input variation occurring due to technical limitations of the system’s stepper motors. The actual “felt” displacement by the spindle depended on a few factors: how far the needle was from a target bundle at the start of manipulation, and how much the whole spindle and cell translated.

### Analysis of live cell manipulations

All analysis was performed in Fiji (Schindelin *et al*., 2012). To ensure proper distance measurements and to account for whole spindle translation, we registered spindles on the GFP-α-tubulin channel using the MultiStackReg Fiji plugin using rigid body registration (Thévenaz *et al*., 1998).

- **Objectively selecting cells for analysis:** Cells were included in the dataset if not negatively affected during the manipulation: cells were excluded if cytokinetic furrowing was visible before the end of manipulation, the spindle collapsed, or if the cells ruptured before finishing manipulation of the midzone bundle.
- **Lateral displacement analysis:** All analysis was performed in a single Z-plane of the GFP-α-tubulin images. Marking X and Y positions of structures was done using the MTrackJ plugin (Meijering *et al*., 2012). The needle position was measured based on the needle fluorescence signal or the GFP-negative shadow of the needle. We defined 2D coordinates for the spindle poles using the GFP-α-tubulin signal, at the center of the brightest signal. A line was drawn along the axis of needle movement and measurements of midzone bundle position (manipulated or neighbors) were determined at the point that they intersected this line for sake of consistency. The perpendicular (shortest) distance between each midzone bundles and the spindle pole-to-pole axis was then measured pre- and post-pull. All bundle movements were normalized to the post-registration distance that the needle moved in that cell. For Fig. 2 D and 2 G, closest bundle to off-target control needle was used as a stand-in for the manipulated bundle in on-target manipulations. For Fig. 2 E and 2 H, only neighbors who showed outward buckling were included; neighbors who started on the same spindle half as the needle and moved away from the direction of needle movement and neighbors who started on the opposite spindle half as the needle and moved in the direction of needle movement were removed (a total of five neighbors for fast pulls and three neighbors for slow pulls). Absolute value of distance traveled from pole-to-pole axis is plotted.
- **Spindle length analysis:** In Fiji the GFP-α-tubulin signal threshold was manually set such that a sharp outline of the spindle was observable. Spindle poles were tracked using the MTrackJ plugin. For consistency, the spindle pole coordinates were marked at the outside edge of the signal apex boundary. Start of manipulation was considered the frame immediately before the computer-programed needle movement began (time = 0 on all time strips) and end of manipulation was considered the first frame after needle movement ended. For expected spindle length comparisons (Fig. 4 F and Fig. 5 G), the average spindle elongation was calculated over the five frames (25 s) prior to needle movement and extrapolated to the frames while the needle was moving.

### Statistics

Data are expressed as mean ± standard deviation (SD) or standard error of mean (SEM) as noted in Figure legends. P-values were calculated by non-parametric Mann Whitney U t-test, Wilcoxon Signed Rank Test, or Wilcoxon matched-pair signed rank test with GraphPad Prism Software, as indicated in Figure legends. We used p < 0.05 as the threshold for statistical significance. Correlations in Fig. 2 E and 2 H were computed using an exponential decay curve:

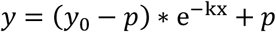

where *y*_0_ is the y value when x (time) is zero, *p* is the *y* value at infinite times, and k is the rate constant.

### Image presentation

Images and movies were formatted using FIJI software. All images display single Z-planes except Fig. 5 A and B which display max intensity projections.

## Supplementary and video legends

S1. **Midzone bundle lateral movement additional analysis. (A)** Natural log transformation of data shown in Fig. 2 C and Fig. 2 F with simple linear regression plotted. Slopes are not different (p = 0.93, analysis of covariance).

Video 1. **Movie of representative on-target fast manipulation of the anaphase spindle**. Fluorescence confocal microscopy movie showing microneedle (BSA Alexa Fluor 647, magenta) manipulation of PtK2 GFP-α-tubulin (green) anaphase midzone bundle. Time (in seconds) is shown where time = 0 is the start of manipulation (7fps). Scale bar = 5 µm. See also Fig. 1D.

Video 2. **Movie of representative off-target fast control manipulation next to the anaphase spindle**. Fluorescence confocal microscopy movie showing microneedle (BSA Alexa Fluor 647, magenta) control manipulation just outside the spindle region in a PtK2 GFP-α-tubulin (green) cell in anaphase. Time (in seconds) is shown where time = 0 is the start of manipulation (7fps). Scale bar = 5 µm. See also Fig. 1E.

Video 3. **Movie of a representative unperturbed anaphase cell**. Fluorescence confocal microscopy movie showing an unperturbed PtK2 GFP-α-tubulin (green) anaphase cell. Time (in seconds) is shown where time = 0 is the start of imaging (7fps). Scale bar = 5 µm. See also Fig. 1F.

Video 4. **Movie of a representative on-target slow manipulation showing lateral midzone bundle coupling in the anaphase spindle**. Fluorescence confocal microscopy movie showing microneedle (BSA Alexa Fluor 647, magenta) manipulation of PtK2 GFP-α-tubulin (green) anaphase midzone bundle and concordant neighbor bundle movement. Time (in seconds) is shown where time = 0 is the start of manipulation (7fps). Scale bar = 5 µm. See also Fig. 2B.

Video 5. **Movie of a representative on-target fast manipulation of the anaphase spindle showing spindle length shrinkage upon force application**. Fluorescence confocal microscopy movie showing fast microneedle (BSA Alexa Fluor 647, magenta) manipulation of PtK2 GFP-α-tubulin (green) anaphase midzone bundle and spindle length shrinkage. White bars mark pre-pull spindle length. Frames during manipulation duplicated for emphasis. Time (in seconds) is shown where time = 0 is the start of manipulation (7fps). Yellow scale bar = 5 µm. See also Fig. 3C.

Video 6. **Movie of a representative slow microneedle manipulation of the anaphase spindle**. Fluorescence confocal microscopy movie showing slow microneedle (BSA Alexa Fluor 647, magenta) manipulation of PtK2 GFP-α-tubulin (green) anaphase midzone bundle. Frames during manipulation duplicated for emphasis. Time (in seconds) is shown where time = 0 is the start of manipulation (7fps). Yellow scale bar = 5 µm. See also Fig. 4A.

Video 7. **Movie of representative on-target fast manipulation of the anaphase spindle with PRC1 siRNA**. Fluorescence confocal microscopy movie showing microneedle (BSA Alexa Fluor 647, magenta) manipulation of PtK2 GFP-α-tubulin (green) anaphase cell treated with PRC1 siRNA. Frames during manipulation duplicated for emphasis. Time (in seconds) is shown where time = 0 is the start of manipulation (7fps). Yellow scale bar = 5 µm. See also Fig. 5D.

---

**Figure S1:**
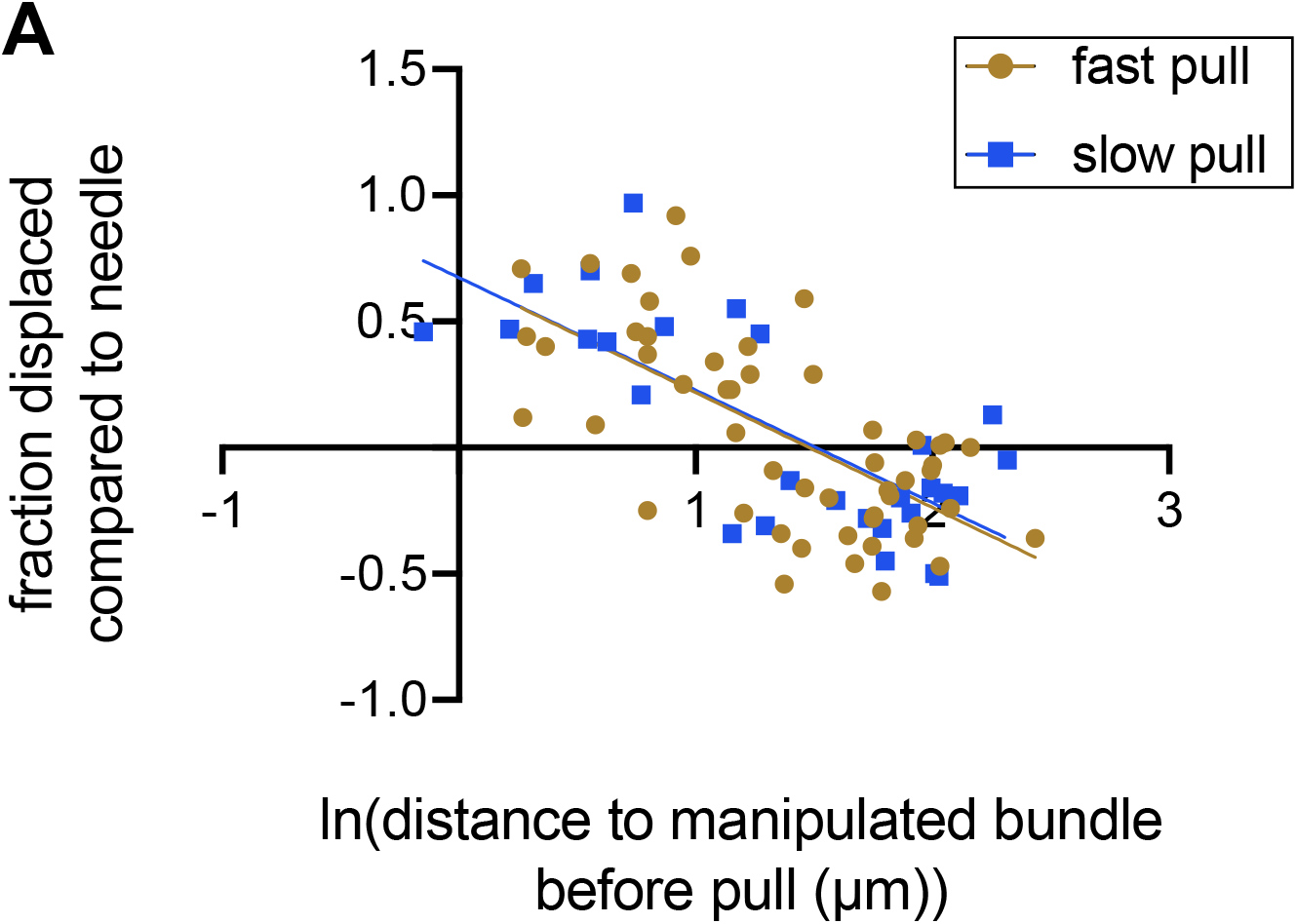
Fast and slow lateral connections show similar trends across space.

